# GAUSS-EM: Guided accumulation of ultrathin serial sections with a static magnetic field for volume electron microscopy

**DOI:** 10.1101/2023.11.13.566828

**Authors:** Kara A Fulton, Paul V Watkins, Kevin L Briggman

## Abstract

Serial sectioning electron microscopy of millimeter-scale 3D anatomical volumes requires the collection of thousands of ultrathin sections. Here we report a high-throughput automated approach, GAUSS-EM, utilizing a static magnetic field to collect and densely pack thousands of sections onto individual silicon wafers. The method is capable of sectioning hundreds of microns of tissue per day at section thicknesses down to 35 nm. Relative to other automated volume electron microscopy approaches, GAUSS-EM democratizes the ability to collect large 3D EM volumes because it is simple and inexpensive to implement. We present two exemplar EM volumes of a zebrafish eye and mouse olfactory bulb collected with the method.

## Introduction

The collection of volumetric electron microscopy data has benefited from several forms of automation^1-7^. These advances can be subdivided into block-face methods that serially ablate tissue within the vacuum chamber of a scanning electron microscope (SBFSEM, FIBSEM, BIBSEM)^1,4,5,7,8^ and serial sectioning methods in which ultrathin sections are first collected and then imaged post hoc (ATUM^3^, MagC^6^). Block-face methods can ablate tissue down to a few nanometers, allowing isotropic resolution in the lateral and axial dimensions, but destroy the sample during acquisition and require specialized microtomes^1^ or ion beams^4,5,7,8^ to be integrated into SEMs. Serial sectioning methods, on the other hand, are limited in minimal section thickness to approximately 30-50 nm^9^, but benefit from a decoupling of the sectioning and imaging phases of data acquisition. That is, after sectioning, section quality can be assessed before a decision is made to proceed with imaging a specimen.

While serial sectioning has been performed by manual ultramicrotomy for decades^10,11^, ATUM and MagC were introduced to automate the collection of sections directly onto conducting substrates. ATUM incorporates a conveyor belt-like pickup system to collect sections onto expensive conductive tape that is subsequently assembled on silicon wafers^3^. An alternative approach, MagC, mitigates the manual wafer assembly of ATUM and increases the packing density of sections on silicon wafers by utilizing a moving magnet to collect sections containing superparamagnetic nanoparticles^6^. However, several limitations remain with this method. Magnetic particles were mixed at a low concentration in a resin and glued onto a tissue sample block, which can in practice lead to a separation of the particles from the section and potential section loss. Sectioning at thicknesses down to 35 nm, a thickness typically required for accurate dense reconstruction in connectomics^12^, has also not been reported with MagC, nor for series of more than a few hundred sections. Finally, like ATUM, the use of motorized actuators leads to an increased complexity and cost of customizing commercial ultramicrotomes.

We sought to improve upon the MagC method to enable the collection of the thousands of 35 nm sections required to scale up to millimeter-scale anatomical volumes by optimizing sample preparation, device design, and automation. Our approach, GAUSS-EM (Guided Accumulation of Ultrathin Serial Sections), uses a static magnetic field to collect sections containing iron oxide nanoparticles onto silicon wafers. Like MagC, this method reduces consumable expenses compared to conductive tapes used in ATUM^13^ and increases the packing density of sections nearly ten-fold. The major advances over MagC are an improved method for dispersing magnetic nanoparticles in resin, the use of a static magnetic field below a collection boat, the demonstration of continuous serial sectioning at 35 nm, and the use of the tissue ultrastructure itself to recover the correct ordering of sections. Our approach enables the collection of large volumes of ultrathin sections with minimal manual intervention at 3-4 times faster sectioning speeds than those previously reported^6,14^ and at a substantially reduced cost.

## Results

We first developed a method to disperse iron oxide particles at a high concentration in the same epoxy resin in which tissue samples were embedded to avoid an interface between two different resins as in MagC. We found that both mechanical mixing and bath sonication were insufficient to disperse the particles, but the use of a probe sonicator in which heat was dissipated during mixing was able to disperse the particles up to a concentration of 30% (w/w) in resin within 30 minutes (Figure 1a, Supplemental Figure 1). The iron/resin mixture was not monodisperse but contained clusters of iron oxide approximately 1 µm in diameter. The mixture was then deposited into a cavity next to a previously embedded tissue sample and polymerized (Figure 1b). The iron concentration and the cross-sectional area of iron/resin exposed when trimming the sample block face were optimized such that 35 nm sections, our target section thickness for connectomic reconstruction, were passively pulled away from the edge of diamond knife beneath a neodymium magnet suspended above the knife boat. Importantly, we found that the 30% concentration of iron nanoparticles was necessary to enable the passive collection of sections and avoid the need for a moving magnet to collect sections as in MagC. We typically form a hexagonal block face that includes a 250 µm wide region of iron/resin oriented to the right of 500-1000 µm wide tissue samples, leading to an iron:tissue block-face ratio substantially below the 50:50 ratio reported for MagC^6^ (Figure 1c).

**Figure 1:**
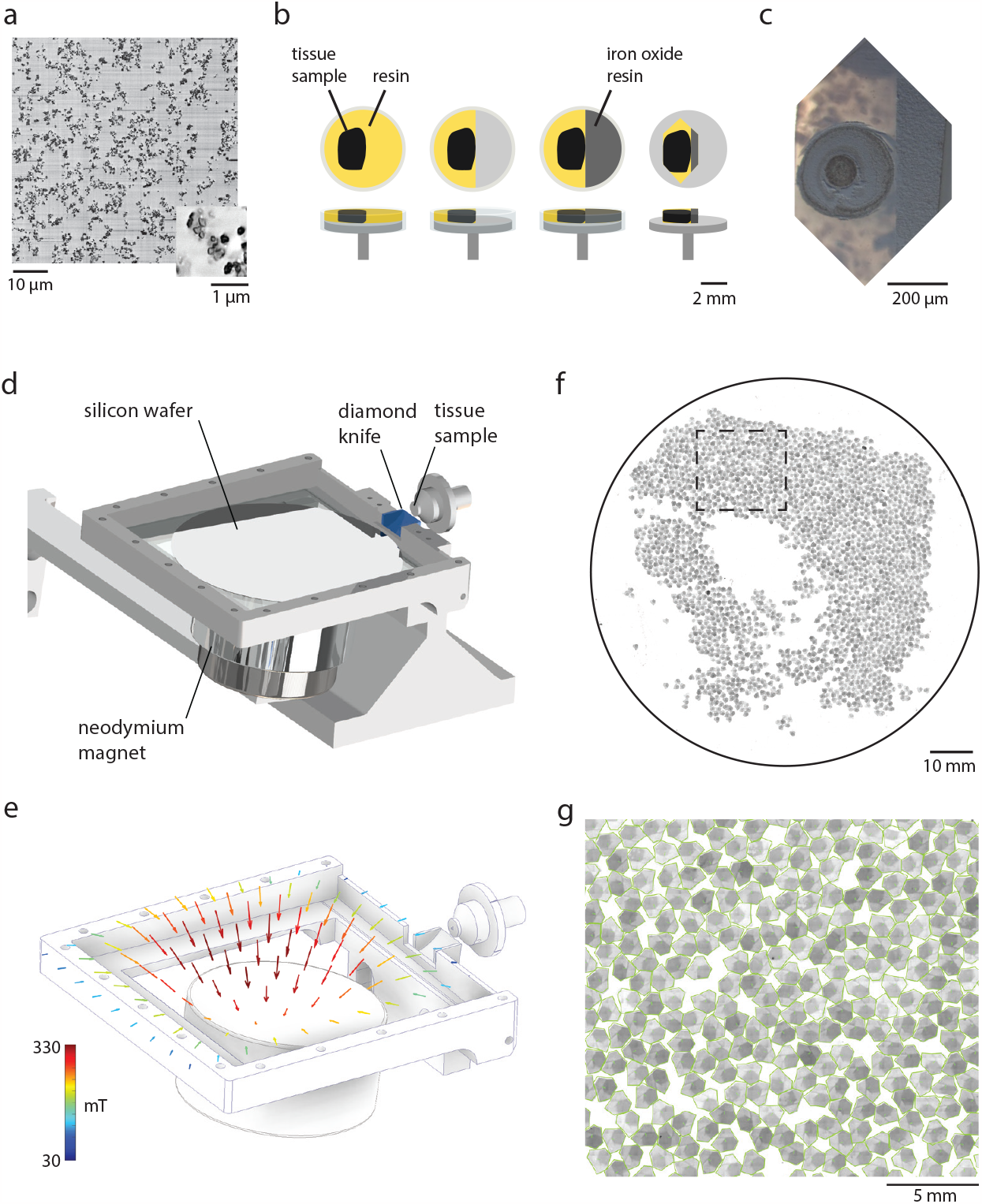
Guided accumulation of ultrathin serial sections with a static magnetic field. **(a)** Electron micrograph of 30% iron oxide dispersed within resin. Inset illustrates iron nanoparticle clusters. **(b)** Sequence of steps to adhere iron/resin mixture to tissue samples. **(c)** Trimmed block face containing a tissue sample and iron/resin mixture. **(d)** Configuration 1 with a custom collection boat for 100 mm silicon wafers and a cylindrical neodymium magnet. **(e)** Magnetic field strengths at the surface of the boat. **(f)** Representative image of 35 nm serial sections collected on a silicon wafer and a magnified view **(g)**.

We next explored two configurations to collect sections with a static magnetic field (see Supplemental Data Files), one in which a cylindrical magnet was positioned below a custom boat (configuration 1, Figure 1d) or in which a spherical magnet was suspended above a boat (configuration 2, Supplemental Figure 2a). For repeatable positioning of the magnets, we quantified the magnetic field strength distribution at the boat surfaces (Figure 1e, Supplemental Figure 2b). For both configurations, a hydrophilized silicon wafer was submerged in the water prior to sectioning on a downward slope oriented towards the front of the boat. During cutting, sections floated to the region of highest magnetic field strength and remained suspended in position. After cutting, water was withdrawn from the boat as sections were held in place by the magnetic field until deposition on the wafer (Figure 2a, Supplemental Video 1). The magnetic field was necessary to hold the sections in place; in the absence of the field, sections dispersed when the water was withdrawn (Figure 2a). For shorter series of sections (<1000), configuration 2 is preferred because the spherical magnet can be positioned close to the diamond knife edge, leading to a stronger pull of sections. A limitation of this configuration is that the magnet obscures the view of sections and a mirror is required to visualize sections from below (Supplemental Figure 2a).

**Figure 2:**
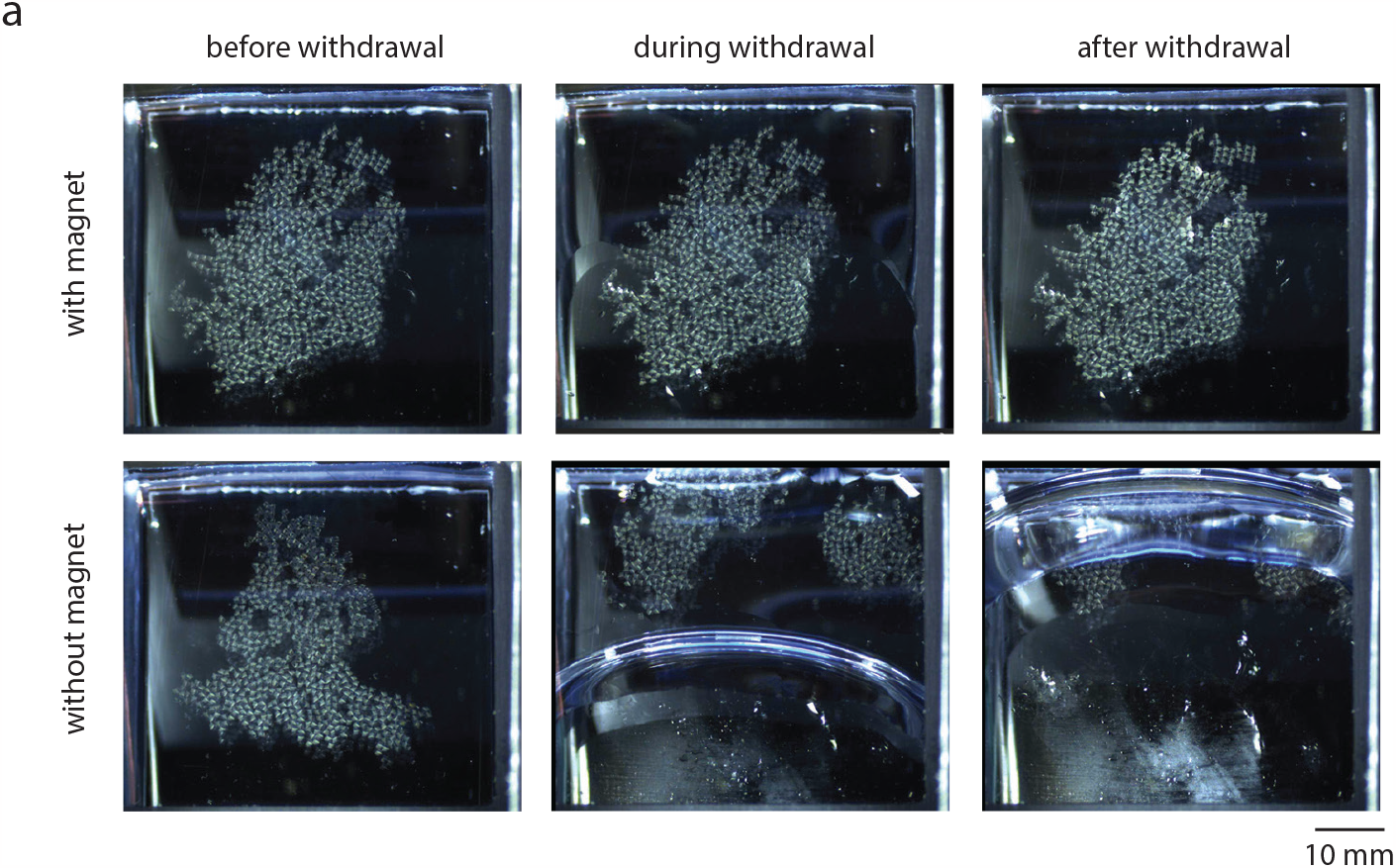
Collection of sections onto silicon wafers. **(a)** Illustration of the location of sections before, during and after the withdrawal of water from the boat both in the presence of the magnet above the boat (top) and absence of the magnet (bottom). For this example, sections were collected on an ITO-coated glass wafer instead of a silicon wafer to visualize the effect of the magnetic field during water withdrawal with a camera from below.

We prefer configuration 1 for longer series of sections (>1000) because the use of a larger 100 mm diameter wafer allows thousands of sections to be densely packed onto a wafer and offers an unobstructed view of the sections during collection. An additional benefit of configuration 1 is that the surface of the boat is covered with a transparent sheet of plastic during sectioning to limit evaporation of water from the boat. Because the size of the magnet restricts how close it can be positioned to the knife edge, we added a glass capillary that delivers a puff of air near the knife edge following each cut (see Materials and Methods). The number of sections that can fit onto a 100 mm wafer depends on the section size, but in practice we typically collect 2000-3000 sections on each wafer (Figure 1f,g). We routinely section at 0.8 – 1.2 mm/s yielding a net collection rate of >1000 sections per hour for block faces of ∼1.5 mm in length.

The sequence in which sections were cut is not preserved once they float onto the water surface, therefore the correct ordering must be determined to assemble a three-dimensional volume. Sections could in principle be tracked by video recording during collection, but we opted for an algorithmic method to solve for the correct ordering of sections following SEM imaging. A SIFT feature^15^ matching algorithm was applied to regions containing tissue for every pairwise combination of 2D-stitched SEM micrographs to assemble a distance matrix among all sections on an individual wafer (Supplemental Figure 3a). We then found the shortest path through this matrix using a traveling salesman problem (TSP) solver (Supplemental Figure 3b, see Code Availability). Sections that do not contain a sufficient number of matching features for the TSP solving step can be semi-automatically placed in the correct sequence (Supplemental Figure 3c). This is typically only required if the imaging contrast is significantly different than most other sections or if a section was damaged during cutting. To assay the robustness of the algorithm, we randomly removed either 50% or 90% of sections from a sequence and re-solved the orderings (Supplemental Figure 3d). In both cases the correct ground-truth ordering was still recovered, except for two swapped sections that needed to be manually corrected when 50% of all sections were randomly removed. Given that missing such a high fraction of sections would be unlikely to yield a useful 3D EM volume anyway, we consider the order solving to be robust to missing sections. We note that a further advantage of this pipeline compared to MagC is the use of the tissue ultrastructure itself to solve the section order and does not require the addition of fluorescent fiducial markers.

**Figure 3:**
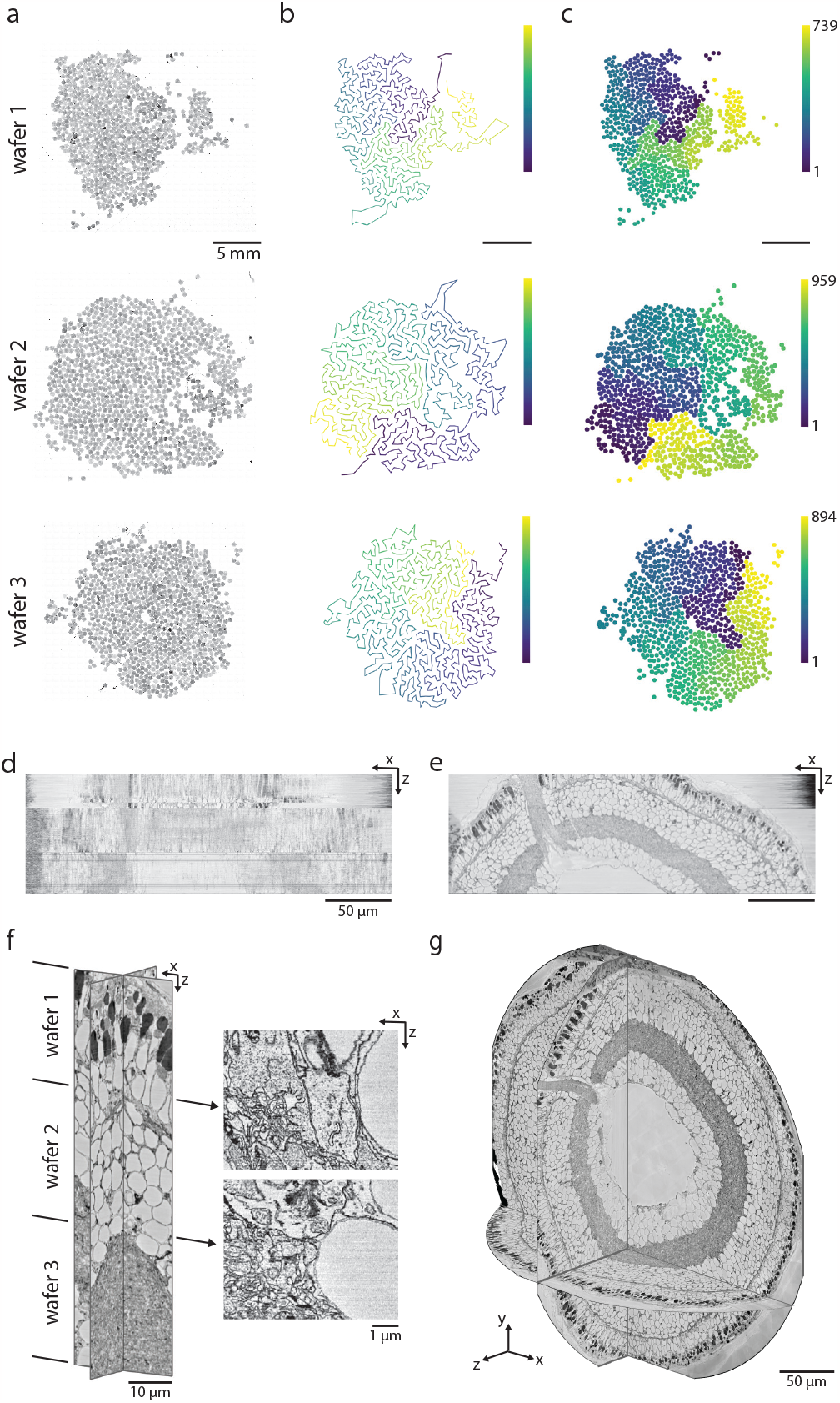
Assembly of sections into 3D volumes. **(a)** Three wafers containing 35 nm sections from a larval zebrafish retina collected with configuration 2. **(b)** Sequence in which sections were imaged. **(c)** Color-coded order of the solved sequence of sections. **(d)** XZ reslice of sections in the imaging order of panel b. **(e)** XZ reslice of sections in the solved order of panel c. **(f)** Magnified XZ reslices, illustrating the transition between wafers 1 and 2 and wafers 2 and 3. **(g)** 3D view of the assembled zebrafish larval retina.

As proof of principle, we collected 3D volumes of a larval zebrafish retina (collected using configuration 2, Figure 3) and from a mouse olfactory bulb (collected with configuration 1, Figure 4). The zebrafish retina (2,592 sections) was collected on three wafer pieces (Figure 3a, left), imaged in an order to minimize SEM stage movements (Figure 3a, middle), and then the section sequence was solved (Figure 3a, right). A XZ virtual slice through the assembled image stack illustrates the imaging order compared to the solved order (Figure 3b, Supplemental Video 2). The olfactory bulb volume (7,495 sections) was collected on four silicon wafers (Figure 4a). To assess the quality of the volumes, we focused on the transitions between wafers and did not observe any gap in the continuity of neurites (Figure 3c, Figure 4c). The final aligned volumes (Figure 3d, Figure 4b) are publicly accessible (see Data Availability).

**Figure 4:**
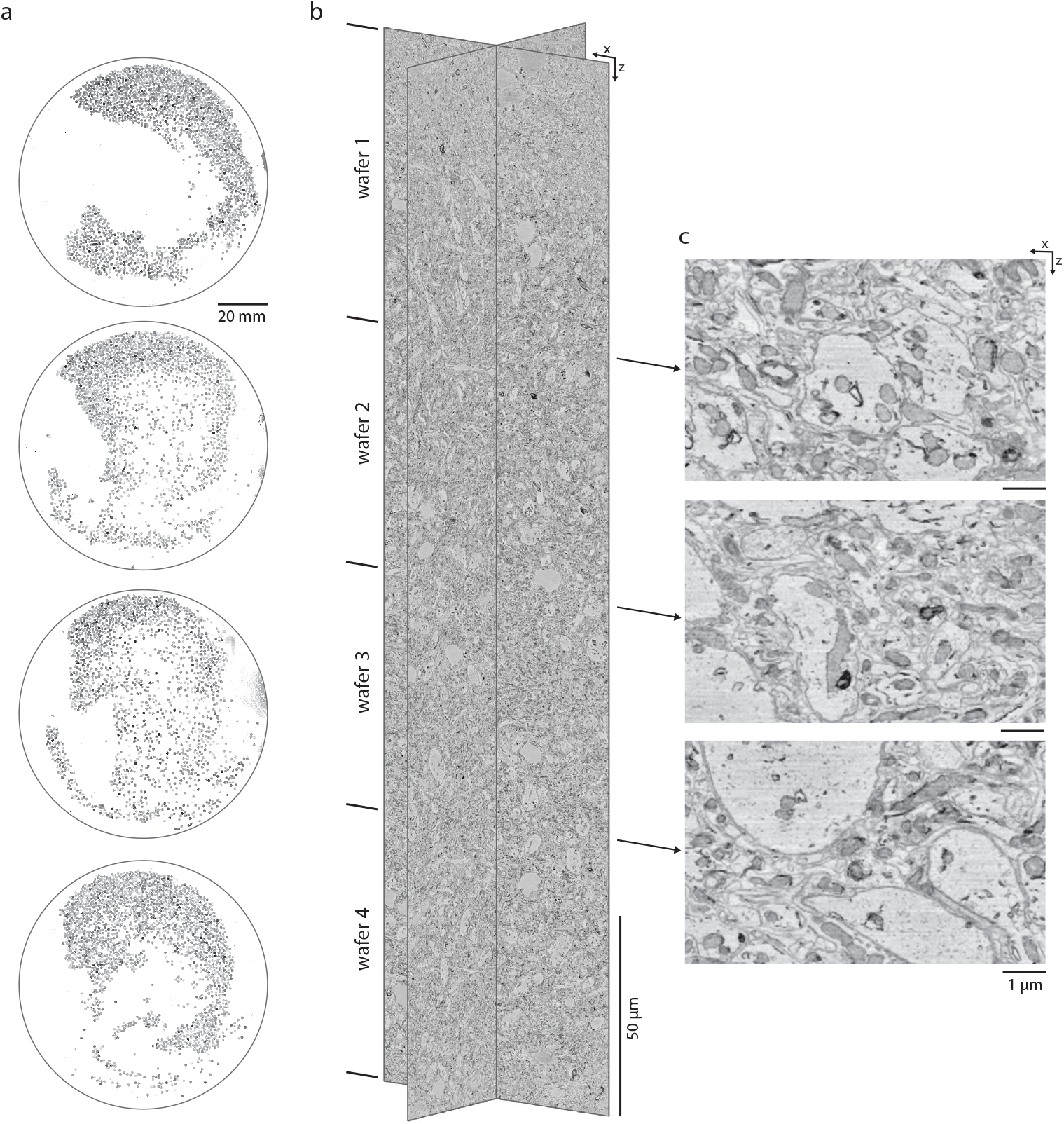
An example volume from the mouse olfactory bulb. **(a)** Light microscope images of sections collected on four 100 mm silicon wafers. **(b)** XZ and YZ reslices through the aligned volume with the boundaries between wafers indicated. **(c)** Higher magnification XZ reslices highlighting the transition between the four wafers in the aligned volume.

## Discussion

Overall, GAUSS-EM is the first ultramicrotomy method that automates the collection of thousands of serial sections by a passive mechanism - a static magnetic field. We routinely cut 35 nm sections at speeds that yield hundreds of microns of tissue cut within a single day. Because the sectioning and imaging steps are decoupled, this method allows one to potentially collect sections at one institution and then image wafers at EM facilities in which high-speed SEMs^16^ are available. The simplicity of GAUSS-EM should allow any laboratory with access to an ultramicrotome to inexpensively implement the method.

In addition to the high-throughput sectioning afforded by GAUSS-EM, the deposition of sections directly onto flat silicon wafers, compared to plastic tapes as in ATUM, allows for a reduction in the imaging overhead caused by autofocusing and autostigmation during SEM acquisition. We typically perform just one round of autofocusing and autostigmation per section, instead of the multiple rounds needed for sections mounted on tapes. To add additional information to EM volumes, GAUSS-EM can be readily combined with correlative light microscopy techniques such as pre- and post-embedding immunohistochemistry^17,18^. Finally, we note that GAUSS-EM is also compatible with hybrid imaging methods in which thicker (>100 nm) sections are collected onto wafers and subsequently milled with an ion beam^4,7^.

## Supporting information

Configuration 1 Parts

Configuration 1 Parts

## Supplemental Figure Legends

**Supplemental Figure 1:**
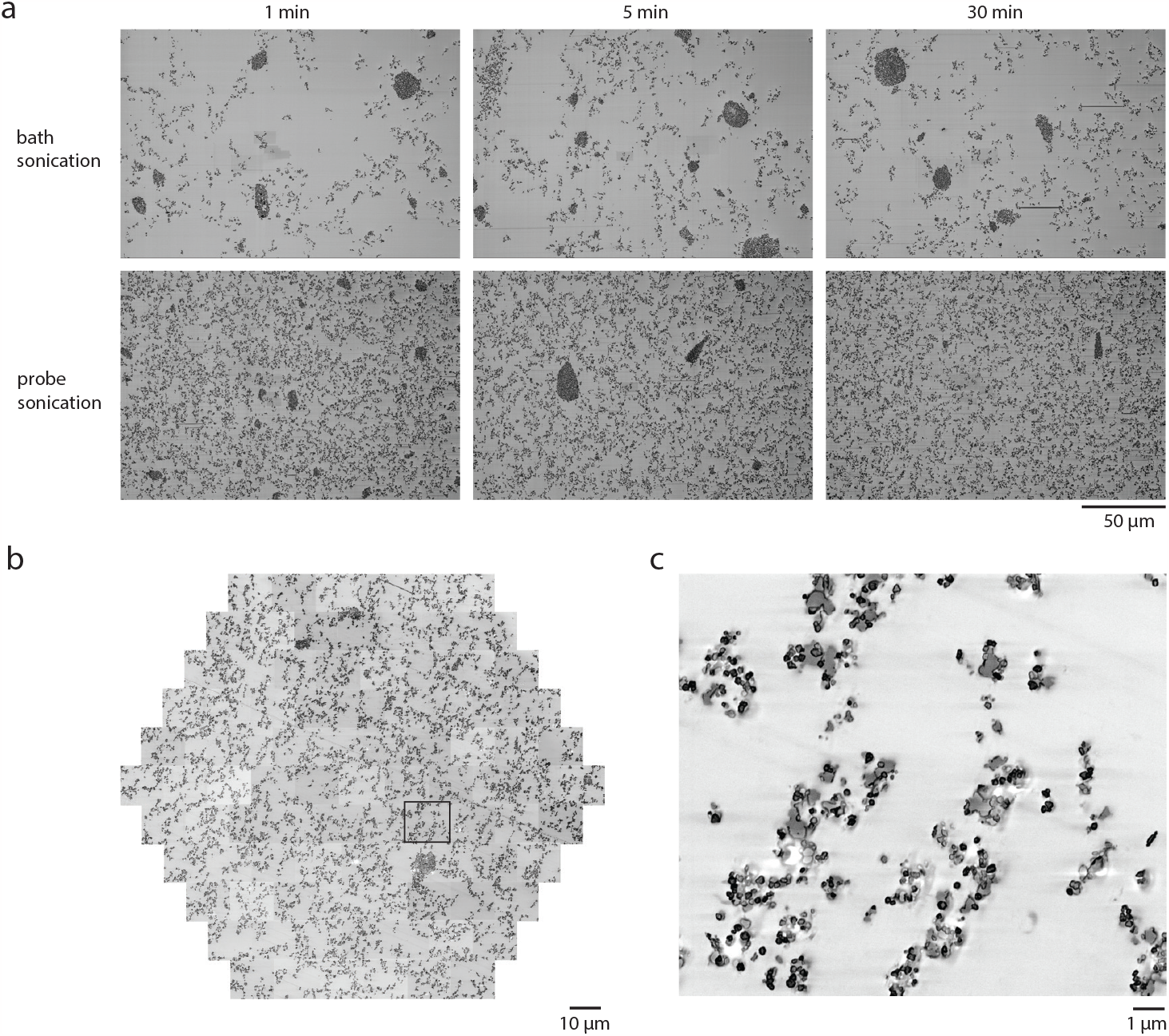
Dispersion of iron oxide in epoxy resin. **(a)** Electron micrographs of 50 nm thick sections taken from samples in which bath sonication (upper row) or probe sonication (lower row) was used for different durations to disperse iron oxide. **(b)** mSEM image of 30% iron oxide dispersed in medium hard Epon. **(c)** Higher magnification of highlighted tile in panel b.

**Supplemental Figure 2:**
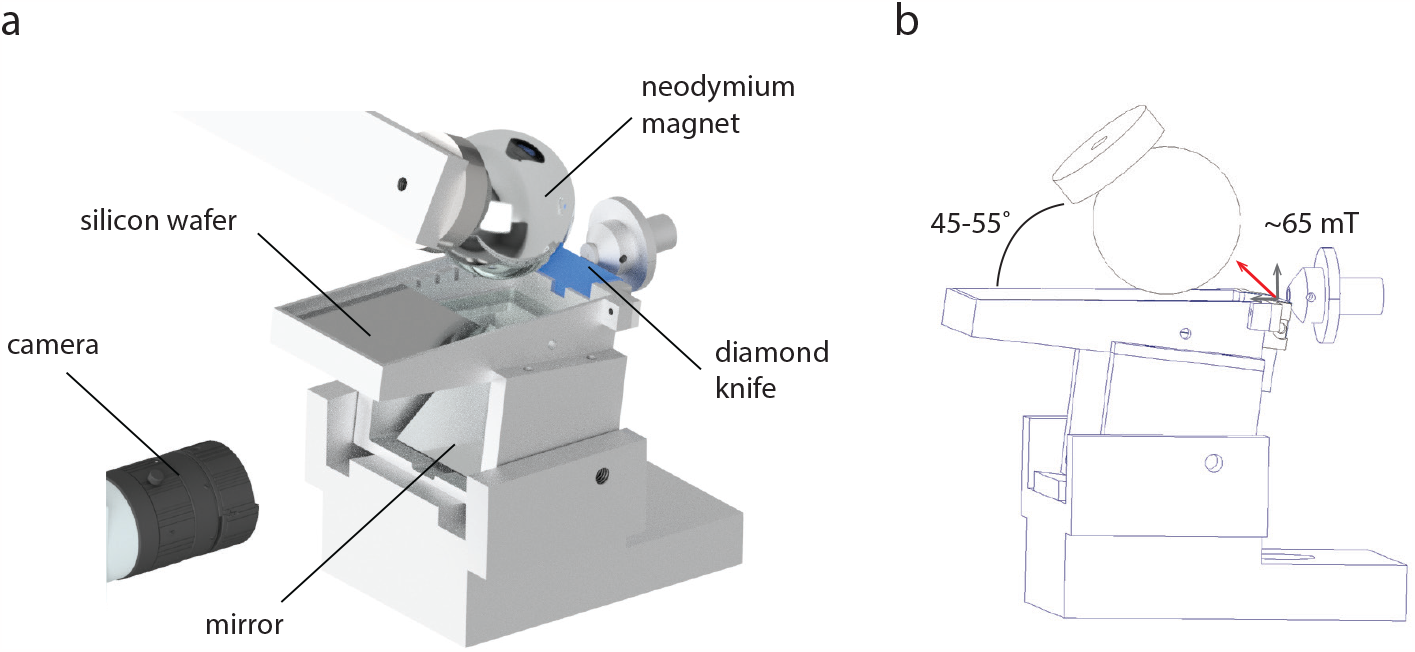
Configuration two for collecting sections. **(a)** Illustration of configuration 2 in which a spherical neodymium magnet is positioned above a custom collection boat designed to accommodate 39 x 42 mm silicon wafers. **(b)** Optimal magnet angle and magnetic field strength measured above the diamond knife edge.

**Supplemental Figure 3:**
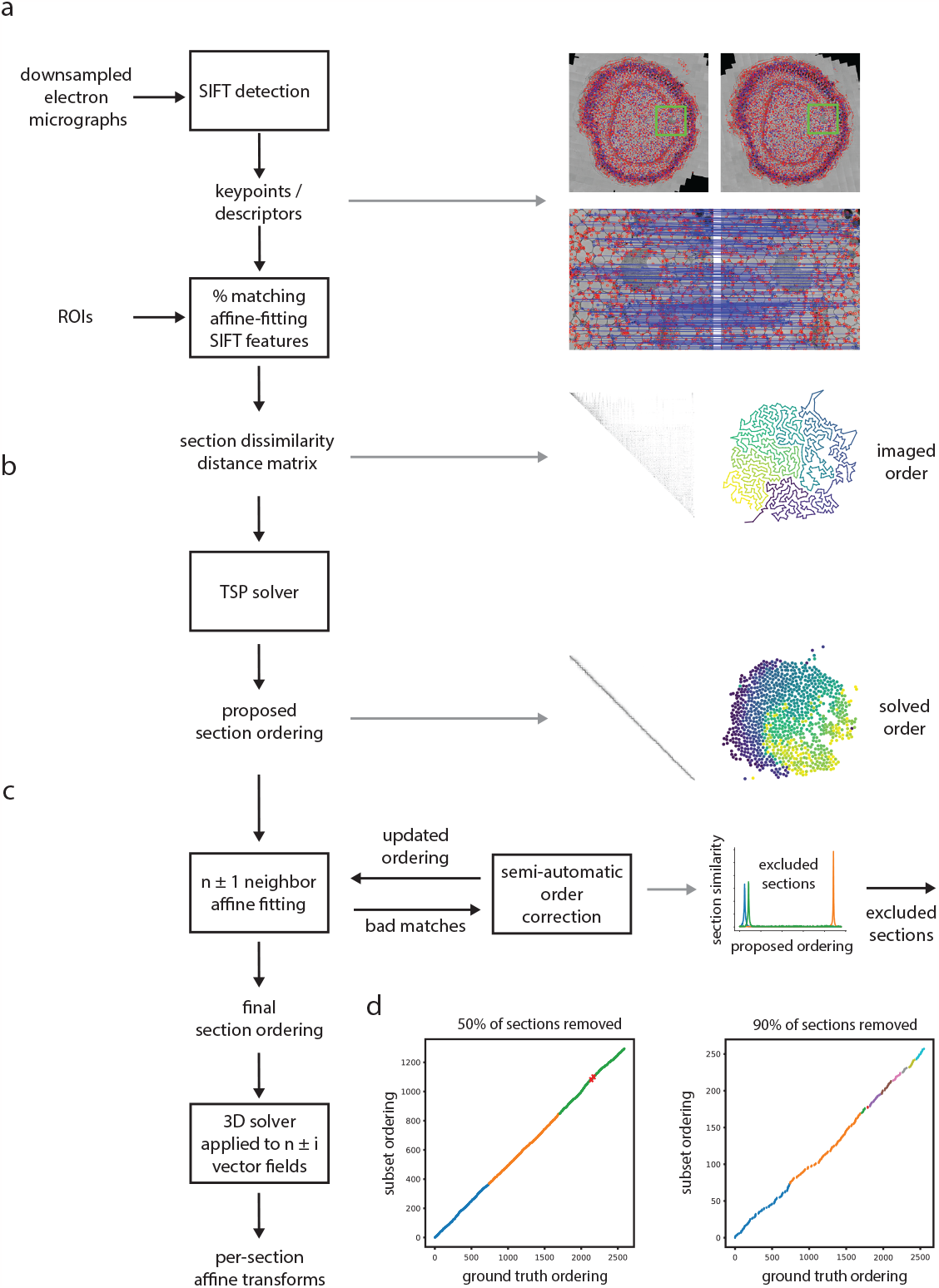
Computational pipeline to solve the order of sections. **(a)** 2D electron micrographs were preprocessed and SIFT features measured from ROIs within each tissue section. Red points indicate detected SIFT features, blue dots and lines indicate matching SIFT features between two sections. A distance matrix was formed among all sections mounted on each wafer using the percent of matching SIFT features as a metric. **(b)** An initial ordering was proposed using a TSP solver to find the shortest path through the distance matrix. Right panels reproduced from Figure 3b,c. **(c)** An affine fitting procedure was used to evaluate the proposed order. Any poorly matched sections were semi-automatically placed in the ordering by finding the location of maximum similarity within the proposed ordering. Example shown of placing 3 slices (blue, green, and orange) within the ordering. During this process, sections that are not sufficiently similar to any sections in the proposed ordering can be permanently excluded. The final section ordering was then aligned with a 3D solver to generate a final affine transform per section. **(d)** 50% (left panel) or 90% (right panel) of sections from the zebrafish retina volume were randomly removed and the order solving was repeated for each wafer. The ordering of the remaining sections was in agreement with the original (ground truth) ordering for each wafer, except for two swapped sections (red X’s). Colored points indicate the solved segments for each wafer.

**Supplemental Figure 4:**
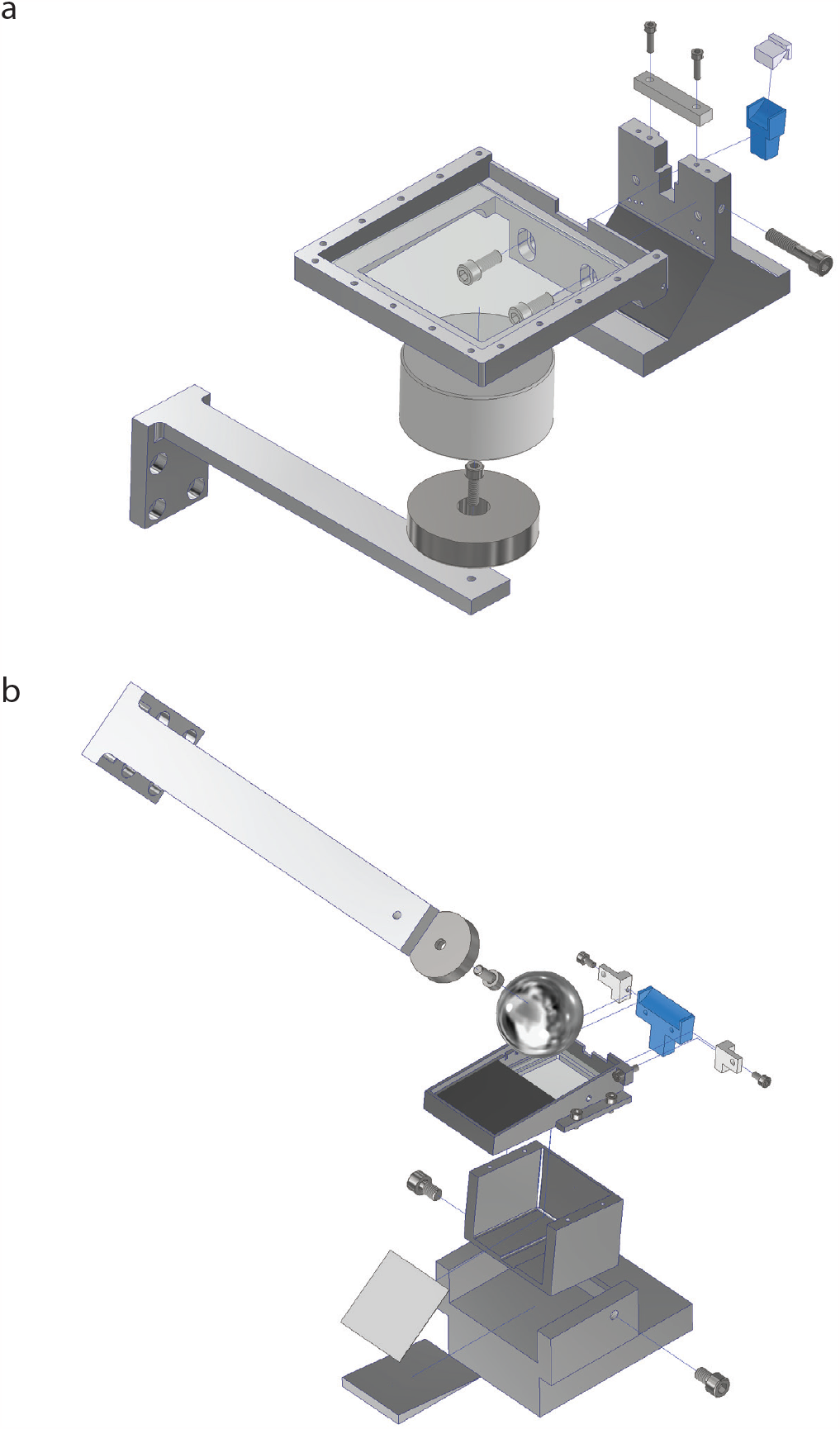
Assembly of custom collection boats. **(a)** Drawing of the assembly of the configuration 1 collection boat. **(b)** Drawing of the assembly of the configuration 2 collection boat.

## Supplemental Data Files

Mechanical part files in STEP format for configuration one and two (Configuration1_Parts.zip; Configuration2_Parts.zip).

Supplemental_Video1.mp4: Representative movie of sections collection using configuration 2. Compares the effect of the magnetic field during withdrawal of water from the boat.

Supplemental_Video2.mp4: Comparison of sections in the order of imaging versus following order solving and alignment.

## Acknowledgements

We would like to thank I Guegel, S Haverkamp, M Pallotto, L Tegethoff and T Yoshimatsu for assistance with sample preparation. We also thank S Irsen for assistance with mSEM imaging and the MPINB mechanical workshop for machining of components. Funding provided by the Max Planck Society.

## Author Contributions

K.A.F and K.L.B developed the method and collected the EM datasets, P.V.W. developed the order solving pipeline. All authors contributed to the writing of the manuscript.

## Materials and Methods

### Animal experiments

All animal experiments were conducted in accordance with the animal welfare guidelines of the Max Planck Society and with animal experimentation approval granted by the Landesamt für Natur, Umwelt und Verbraucherschutz Nordrhein-Westfalen, Germany.

An adult (C57BL/6) mouse was first anesthetized with isofluorane before swift decapitation. The brain was carefully removed from the skull, and 300 µm horizontal sections from the olfactory bulb were cut on a vibratome (Leica) and briefly stored in a cold carboxygenated (95% O_2_/5% CO_2_) ACSF solution (300– 320 mOsm) containing (in mM): 124 NaCl, 3 KCl, 1.3 MgSO_4_.7H_2_O, 26 NaHCO_3_, 1.25 NaH_2_PO_4_.H_2_O, 20 glucose, 2 CaCl_2_.2H_2_O. Sections were then immersion-fixed in 4% paraformaldehyde (Electron Microscopy Sciences) and 2% glutaraldehyde (Electron Microscopy Sciences) using a protocol to preserve extracellular space^19^.

A 6 dpf larval zebrafish was anesthetized in 0.01% tricaine, the eyes enucleated, and immersion fixed in 2% glutaraldehyde in 150 mM cacodylate overnight.

### EM staining and resin embedding

The samples were stained as previously described^20^. Briefly, the samples were stained in a solution containing 2% osmium tetroxide, 3% potassium ferrocyanide, and 2mM CaCl_2_ in 150 mM CB for 2 hrs at 4° C, followed by 1% thiocarbohydrazide (1 hr at 50° C), and 2% osmium tetroxide (1 hr at room temperature). The samples were then stained with 1% aqueous uranyl acetate for 6 hrs at 45° C and lead aspartate for 6 hrs at 45° C. The tissue was dehydrated at 4° C through an ethanol series (70%, 90%, 100%), transferred to propylene oxide, infiltrated at room temperature with 50%/50% propylene oxide/Epon, and then 100% Epon. Both samples were embedded in medium hard Epon^21^ (14120; Electron Microscopy Sciences) and cured on aluminum stubs (75638-10; Electron Microscopy Sciences) at 60° C for 24 h.

### Iron/resin preparation

We tested several iron oxide nanoparticles for their ability to disperse in epoxy resin and the strength of the magnetic pull when sectioned at 35 nm. The optimal formulation was iron oxide II,III nanopowder (50-100 nm size particles; #637106; Sigma-Aldrich). 10 mL of medium hard Epon was prepared in a 20 mL glass scintillation vial by weight but without the addition of the BDMA accelerator and mechanically swirled until evenly mixed. The mixture was warmed in a 60° C oven for 15 minutes to reduce viscosity and 30% weight/weight iron oxide was added to the Epon mixture and vortexed for 1 minute. Using a 450 W digital probe sonicator (Branson W-450 D), the mixture was then sonicated at 20% amplitude for 30 minutes in 5 minute intervals with the sonicator probe fully immersed in the scintillation vial. To dissipate heat during sonication the scintillation vial was surrounded in a container with ice cold water. Following sonication, the accelerator was added and mechanically swirled. We observed equivalent dispersion in other embedding resins including different hardness formulations of Epon as well as Durcupan and Spurr’s resins.

### Sample block preparation

To create a cavity for the iron/resin mixture, excess epon was trimmed from one side close to the sample parallel to the cutting direction. The aluminum stub was then surrounded with a tight-fitting thin plastic tubing to create a well. A drop of the freshly prepared iron/resin was then deposited with an insect pin into the cavity. The sample and iron/resin were then cured at 70° C for 24-48 hours. To minimize compression along the cutting direction (section length) and to ensure that sections detach from the knife edge and migrate towards the magnet, we shaped the block with pointed leading and trailing edges. This creates a minimal contact area of each section with the knife edge such that the epon of the previous section does not adhere to the following section or the knife edge. Samples were trimmed with a dry diamond knife to block face sizes approximately 1200-1500 µm long (parallel to the cutting direction) and 750-1000 µm wide including ∼250 µm of the iron/resin to the right of the tissue.

### Assembly of collection boats and sectioning procedure

The custom collection boats were machined from aluminum and consist of two parts, a frontend to clamp a diamond knife and a backend collection boat that is sized for either configuration one or two. To assemble the boats, a diamond knife (35° or 45° Ultra or Ultra Jumbo knives, Diatome) is first clamped into the frontend and held at the manufacturer specified clearance angle (typically 0 degrees or 6 degrees). The knife edge was then covered with a 3D printed cover and secured in place with a clamping bracket. The rear portion of the knife was then milled to a depth flush with the frontend holder. The milling of knives does not preclude the ability to have them resharpened by the manufacturer (Diatome). The backend collection boat was then screwed to the frontend and then interface between the diamond knife and backend was made water-tight by applying a thin bead of cyanoacrylic glue. The bottoms of the backend collection boats were fitted with either plastic or glass and sealed with cyanoacrylic glue. For assembly of the boats see Supplemental Figure 4. All sectioning was performed with a Leica UC7 ultramicrotome.

#### Configuration One

For collection with configuration one, a neodymium pot magnet with counterbore hole (ZTN-32; supermagnete) was screwed to a support arm that is attached to a rotary stage (Thorlabs) and XYZ micrometer positioner (Thorlabs). A 70 mm diameter cylindrical neodymium magnet (S-70-35-N; supermagnete) was then held in place by the attraction to the pot magnet. Care should be taken when handling the magnets due to the high field strength. The rotary stage allows the relative angle of the magnets to be fine-tuned with respect to the bottom of the backend collection boat. To prepare for sectioning, a 100 mm diameter, 300 µm thick silicon wafer (BO14072; Siegert Wafer) was first glow discharged (Q150R ES; EMS) to create a hydrophilic surface. The wafer was placed on the bottom and the boat filled with Millipore deionized water. Control of the water level was accomplished via a side port that allowed water to be perfused or withdrawn using a syringe pump (NE-1000; New Era Pump Systems). For repeatable positioning of the magnet below the collection boat, the field XYZ components of the magnetic field strength were measured in a grid pattern from the surface of the boat using a teslameter magnetometer (Projekt Elektronik Teslameter FM 302). The rate at which sections are drawn toward the backend collection boat depends on the strength of the magnetic field at the knife edge, the section thickness, and the cross-sectional area of iron oxide/resin within each section. To assist sections to move toward the backend and prevent sections from accumulating near the knife edge, an optional air puffer was used. The air puffer consisted of a tapered glass capillary attached to a XYZ translator (Thorlabs) and oriented to puff air at the water surface approximately 1 mm behind the knife edge. This had the effect of drawing sections away from the edge of the knife and pushing them toward the backend collection boat. The air puffer was supplied with house compressed air and was controlled with a solenoid pinch valve (PM-0815W; Takasago Fluidic Systems) that was triggered at the end of each downward swing of the microtome cutting arm. Triggering was achieved by mounting a 3mm infrared beam break sensor (Adafruit) on either side of the microtome cutting arm that was read by a microcontroller (Duo; Arduino), which then generated a trigger signal to the pinch valve on each break of the IR beam.

During sectioning, a plastic barrier was placed atop the backend collection boat to reduce the rate of evaporation from the boat as well as prevent dust from falling onto the water surface. Following sectioning, sections were deposited onto the silicon wafer by withdrawing water from the boat at a rate of 5-10 mL/min with the syringe pump. The wafer was then removed from the boat with plastic forceps and any residual water on the surface was evaporated by placing the wafer on a 60° C peltier heating plate (BSH300; Benchmark Scientific) for a few minutes.

#### Configuration Two

For collection with configuration two, a 32 mm diameter neodymium pot magnet with counterbore hole (ZTN-32; supermagnete) was screwed to a support arm that is attached to a rotary stage (Thorlabs) and XYZ micrometer positioner (Thorlabs). A spherical 40 mm diameter neodymium magnet (K-40-C; supermagnete) was then held in place by the attraction to the pot magnet. Care should be taken when handling the magnets due to the high field strength. The rotary stage allows the relative angle of the magnets to be fine-tuned with respect to the surface of the backend collection boat. To prepare for sectioning a silicon wafer (KristallTechnologie S4974) was cut with a wafer saw to a 39 x 42 mm^2^ rectangle and hydrophilized (PELCO easiGlow) with a negative polarity to air and 20 mA current for 5 minutes. The wafer was placed toward the rear of the backend and the boat was filled with deionized water. Control of the water level was accomplished via a side port that allowed water to be perfused or withdrawn using a syringe pump. For repeatable positioning of the magnet above the collection boat, the field XYZ components of the magnetic field strength were measured above the knife edge using a teslameter magnetometer (Projekt Elektronik Teslameter FM 302). The rate at which sections were drawn toward the backend collection boat depends on the strength of the magnetic field at the knife edge, the section thickness, and the cross-sectional area of iron oxide/resin within each section. For visualization of sections on the water surface during cutting, a USB camera was oriented toward a 45° mirror underneath the boat. When ready to collect sections, the wafer was slid forward underneath the sections and water was withdrawn at a rate of 10 ml/min. The wafer was then removed from the boat with plastic forceps and any residual water on the surface evaporated by placing the wafer on a 60° C peltier heating plate for a few minutes.

### Serial sectioning

The zebrafish eye, stained and embedded as described above, was trimmed to a block face width of 420 µm (including 140 µm of iron oxide/resin) and length of 620 µm. The sample was sectioned with a 35 nm section thickness at a speed of 0.8 mm/s using the configuration 2 collection boat. Three wafers (S4974; KristallTechnologie) cut to 39 x 42 mm squares were collected containing 739, 959, and 894 sections, respectively.

The vibratome section of the mouse olfactory bulb, stained and embedded as described above, was trimmed to a block face width of 1000 µm (including 250 µm of iron oxide/resin) and length of 1500 µm. The sample was sectioned with a 35 nm section thickness at a speed of 1.2 mm/s using the configuration 1 collection boat. Four wafers were collected containing 1983, 1865, 1678, and 1969 sections, respectively.

The presence of iron oxide nanoparticles in the block did not lead to any noticeable damage to diamond knives, as we have used the same diamond knife for multiple large-scale 35 nm serial section experiments. Within an experiment, after every few thousand sections, we move the knife to the right so the left side of the sample block that contains tissue is cut with a fresh knife edge.

### SEM Imaging

Both volumes were imaged using a 91-beam multibeam scanning electron microsope (mSEM; Zeiss) with a 15 µm beam pitch. The mSEM was controlled via the Zeiss mSEM API. Regions of interest were defined with a template matching-based segmentation, similar to WaferMapper^22^, of each section on a wafer in Matlab (Mathworks) and then converted to hexagonal fields of view (mFOVs) using the mSEM API. During SEM imaging, we perform one round of autofocus and autostigmation per section over the iron containing region. Sections were imaged with a 50 ns dwell time, 4 nm pixel size and 1.5 kV landing energy. The zebrafish eye dataset contains (in x,y,z) 67348 x 70125 x 2573 voxels (excluding the surrounding resin) and the mouse olfactory bulb dataset downsampled to 16 nm in x,y contains in (x,y,z) 4000⍰×14000⍰× ⍰7495 voxels.

### Alignment and assembly of 3D EM volumes

#### Preprocessing

2D stitching between individually acquired image tiles (corresponding to individual mSEM beams) was performed by calculating 2D cross correlations between neighboring tiles on the same section. Tile positions were solved for using these translations resulting in a global best fit per section (a least squares solution). 2D-stitched section images were corrected for between-tile gradients or offsets by blending. Images were also normalized between sections for brightness and contrast, because the section order solving is sensitive to these differences.

#### Order solving

2D-stitched images were downsampled (128 nm) and then SIFT features^15^ were detected on each section, with keypoints constrained to the ROI region defined before imaging to eliminate potential spurious descriptor matches from non-tissue containing areas (Supplemental Figure 3a). An image distance metric was calculated between all sections on a single wafer based on the percentage of matching SIFT features. The section order was then resolved by applying an exact traveling salesman problem solver^23^ to this distance matrix, generating an initial proposed ordering (Supplemental Figure 3b). Bad matches in the proposed ordering were detected as sections that did not fit to an affine transformation with their neighbors. These order problems were then resolved semi-manually. For example, sections that did not fit in the proposed ordering were compared again against all sections, but now as a function of this proposed ordering, and then inserted at minimum locations of the distance metric (Supplemental Figure 3c). Any sections that suffer from uncorrectable artifacts (e.g. a thin section substantially less than 35 nm and therefore of insufficient contrast) were excluded from the volume at this step. Once the order was solved, sections were aligned by an iterative 3D alignment pipeline (Watkins, Jelli, and Briggman, under review) similar in strategy to previously described EM alignment pipelines^24^.

To estimate the robustness of the order solving procedure, we randomly removed sections from the zebrafish retina volume and repeated the order solving. For cases in which 50% or 90% of sections were removed, the solved order of the remaining sections for each wafer remained in the correct sequence compared to the ground truth sequence, except for 2 swapped sections with 50% removed (Supplemental Figure 3d). With 90% of sections removed, discontinuous segments within the solved order appeared, but the sequence within these segments was correct compared to the ground truth.

### Cost estimation

The one-time cost to implement GAUSS-EM is on the order of several thousand dollars which includes custom machining of the aluminum collection boats, magnets, syringe pump, microcontroller, pinch valve, hot plate, and teslameter. The consumable costs are on the order of a couple hundred dollars per experiment consisting solely of the cost of silicon wafers (currently ∼$25/wafer), iron oxide nanoparticles and embedding resin. Not included are the costs of common equipment for an electron microscopy facility such as a commercial ultramicrotome, probe sonicator, glow discharger and diamond knives.

## Data availability

The zebrafish retina dataset is viewable at https://webknossos.mpinb.mpg.de/links/4ig-0q1evJ649zfo. The mouse olfactory bulb dataset is viewable at https://webknossos.mpinb.mpg.de/links/2VjYQ1O3vKUhRZId.

## Code availability

An example for the section order solving procedure and source code are available at https://github.com/mpinb/gauss-em

